# Striking Differences in Tetrabromobisphenol A (TBBPA) Deconjugation by Human Fecal Microbiota: The Pilot-10 Human Donor Study

**DOI:** 10.64898/2026.11.13.766573

**Authors:** Cayen Rong, Joe Lim, Ximeng Liu, Lingjun Li, Shiquan Cui, Haiwei Gu, Andrew Annalora, Julia Yue Cui

**Affiliations:** Environmental and Molecular Toxicology, Oregon State University; Institute for Systems Biology, Seattle, WA; College of Health Solutions, Arizona State University, Phoenix, AZ

## Abstract

Tetrabromobisphenol A (TBBPA) is a widely used brominated flame retardant with multiple adverse health effects. Although TBBPA undergoes phase II conjugation to facilitate elimination, gut microbial deconjugation may regenerate unconjugated TBBPA, increasing intestinal re-exposure. However, the extent of interindividual variability and its underlying microbial determinants remain poorly understood. To address this critical knowledge gap, we hypothesized that human fecal microbiota exhibit donor-dependent differences in TBBPA-sulfate (S-TBBPA) deconjugation and potential downstream debromination to tribromobisphenol A (Tri-BBPA). In our Pilot-10 Human Donor Study (PM10), fecal microbiota from five healthy adult male and five female human donors were incubated anaerobically with S-TBBPA at 37°C for six hours. Heat-inactivated microbiota served as controls. S-TBBPA, unconjugated TBBPA, and Tri-BBPA were quantified using targeted liquid chromatography-mass spectrometry (LC-MS). Metagenomic shotgun sequencing (MGS) was performed followed by microbial taxonomic and gene-family profiling using MetaPhlAn v4.11 and HUMAnN v3.9, respectively. Unheated fecal microbiota from all donors regenerated TBBPA, whereas heat inactivation nearly abolished its formation, supporting microbial enzyme-dependent deconjugation of S-TBBPA. The proportion of regenerated TBBPA varied substantially among donors, accounting for approximately 10-50% of the combined detected S-TBBPA and TBBPA. No statistically significant sex-associated differences were detected. Tri-BBPA was undetectable or negligible across incubations. Notably, the cumulative sulfatase-maturation enzyme gene family abundance was positively associated with LC-MS-measured S-TBBPA deconjugation activity, whereas the cumulative arylsulfatase and other sulfatase gene abundances showed no significant associations. Furthermore, 12 taxonomically stratified microbial desulfation-related gene-family annotations exhibited nominally significant positive correlations with deconjugation activity, highlighting potential roles for specific microbial sulfatases and their maturation enzymes in TBBPA reactivation. Collectively, our PM10 pilot study revealed substantial interindividual variability in microbial TBBPA-sulfate deconjugation and identified sulfatase-maturation enzyme gene abundance as a potential indicator of activity. These findings highlighted an underexplored pathway of xenobiotic reactivation and support future personalized microbiome-directed strategies to reduce internal TBBPA exposure and associated health risks.

## Introduction

Tetrabromobisphenol A (TBBPA) is one of the most widely used brominated flame retardants (BFRs), historically accounting for approximately 60% of global BFR use ^1–3^. Its extensive application in electronics, textiles, and plastics has contributed to widespread environmental contamination and ongoing human exposure, primarily through dietary intake and dust ingestion (**Figure 1A**) ^4, 5^. TBBPA has been detected in human serum, breast milk, and urine, while experimental studies have demonstrated multiple toxicological effects, including endocrine disruption, particularly interference with thyroid hormone signaling, as well as neurodevelopmental, immune, reproductive, and hepatic toxicity ^5–7^. Moreover, the International Agency for Research on Cancer (IARC) classified TBBPA as a Group 2A carcinogen (probably carcinogenic to humans) based on sufficient evidence in experimental animals and strong mechanistic evidence ^8^. In summary, given its widespread use, persistent environmental occurrence, ongoing human exposure, and potential adverse health effects, investigating the metabolism and toxicity of TBBPA remains highly relevant and timely for understanding its human health risks and improving environmental health risk assessment.

**Figure 1.**
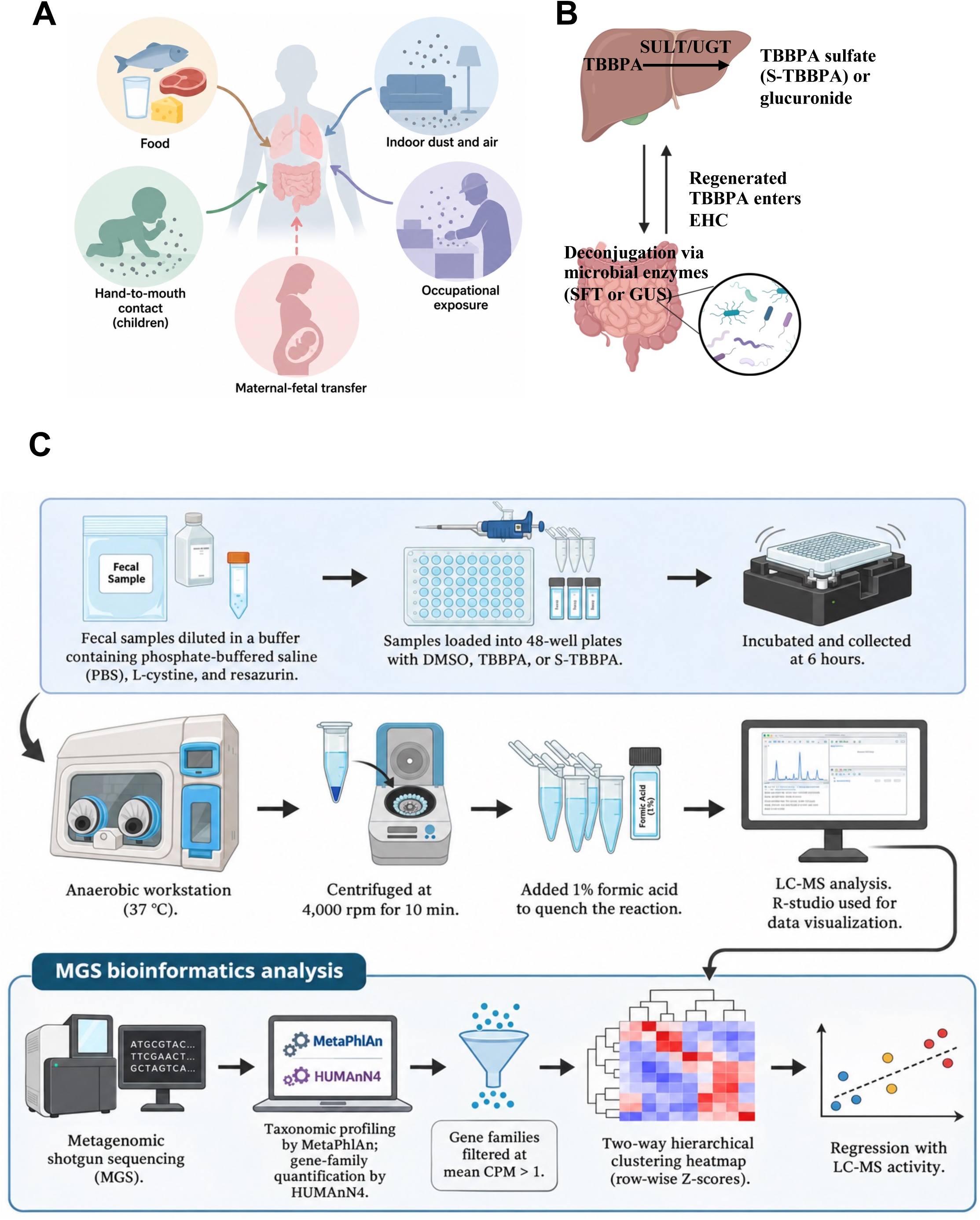
Study rationale and experimental design the PM10 pilot study. **A.** Schematic illustration of human exposure pathways to TBBPA, including dietary intake, indoor dust and air, hand-to-mouth exposure, occupational exposure, and maternal-fetal transfer. The illustration was created using BioRender.com. **B.** Proposed role of the gut microbiome in TBBPA metabolism and enterohepatic recycling. Following exposure, TBBPA undergoes host-mediated phase II conjugation by sulfotransferases (SULTs) and UDP-glucuronosyltransferases (UGTs), generating TBBPA-sulfate and TBBPA-glucuronide, respectively. Conjugated metabolites entering the intestinal tract may undergo microbial deconjugation by sulfatases or β-glucuronidases, regenerating unconjugated TBBPA and potentially facilitating intestinal reabsorption and enterohepatic recirculation. The illustration was created using BioRender.com. **C.** Experimental workflow for assessing microbiome-mediated TBBPA-sulfate deconjugation. Human fecal samples from 10 donors were suspended in phosphate-buffered saline supplemented with L-cysteine and resazurin and incubated with DMSO, TBBPA, or TBBPA-sulfate under anaerobic conditions at 37°C for 6 hours. Samples were centrifuged at 4,000 rpm for 10 minutes, and reactions were quenched with 1% formic acid before LC-MS analysis. Metagenomic shotgun sequencing (MGS) was performed on samples from nine donors using paired-end sequencing (2 × 150 bp; approximately 5 million reads per sample). Microbial taxonomic profiling was conducted using MetaPhlAn v4.11, and gene-family abundances were quantified using HUMAnN v3.9 with UniRef90 annotations. Gene families with mean abundance greater than 1 count per million (CPM) were retained for functional analyses, including two-way hierarchical clustering of row-wise Z-score-normalized abundances and correlation analyses with LC-MS-measured TBBPA-sulfate deconjugation activity. Disclaimer: The experimental workflow illustration in Fig. 1C was created using generative AI and subsequently reviewed and edited by the authors to ensure scientific accuracy. Generative AI was used solely for graphical illustration and was not involved in scientific data generation, analysis, or interpretation.

Following absorption, TBBPA undergoes extensive phase II metabolism, primarily through glucuronidation and sulfation, which facilitate its elimination (**Figure 1B**) ^9, 10^. Animal studies have further demonstrated biliary excretion of TBBPA conjugates and provided evidence suggesting enterohepatic circulation, indicating that these metabolites may enter the intestinal lumen and encounter the gut microbiome ^11^. Gut microorganisms possess diverse deconjugating enzymes capable of reversing host-mediated phase II metabolism, potentially regenerating parent xenobiotics and modifying their intestinal availability, reabsorption, and biological activity.

Although both glucuronidation and sulfation contribute to TBBPA metabolism, glucuronide conjugates are more consistently detected in human plasma ^9, 12^, while little known regarding why TBBPA-sulfate was only detected in some individuals but not all human plasma samples. Thus, the present study specifically focused on TBBPA-sulfate to investigate what contributes to the interindividual variations in the TBBPA-sulfate metabolism. Specifically, we investigated microbial desulfation, which is an understudied mechanism of microbiome-mediated xenobiotic deconjugation (**Figure 1B**). While microbial β-glucuronidase-mediated hydrolysis of xenobiotic glucuronides has been extensively characterized, including its roles in enterohepatic recycling, drug reactivation, and gastrointestinal toxicity ^13, 14^, in contrast, the substrate specificity and toxicological implications of gut microbial sulfatases remain less well understood. Recent studies have demonstrated substantial functional diversity among human gut microbiome-encoded sulfatases, highlighting their potential to hydrolyze sulfated metabolites ^15^. However, whether human gut microorganisms can deconjugate TBBPA-sulfate to regenerate unconjugated TBBPA, and the extent to which this activity varies across individuals, remain largely unexplored.

In addition to TBBPA-sulfate deconjugation, we also investigated the potential formation of tribromobisphenol A (Tri-BBPA) as a marker of subsequent microbial debromination. Previous studies have demonstrated that selected human gut bacterial strains can transform brominated bisphenols through microbial biotransformation pathways, including debromination ^16^. Furthermore, environmental anaerobic microbial communities have been shown to convert TBBPA into less brominated metabolites, including Tri-BBPA, through sequential reductive debromination ^17^. Importantly, Tri-BBPA exhibits greater estrogenic activity than TBBPA in human cell-based assays ^18^ and has been associated with hepatotoxic effects in experimental animals ^19^. However, the extent of the human fecal microbiome community in reductive debromination of TBBPA to Tr-BBPA is not known. Therefore, we also quantified Tri-BBPA alongside TBBPA-sulfate and unconjugated TBBPA in human fecal incubations to assess whether microbial deconjugation was accompanied by debromination.

Given the substantial interindividual variability in gut microbial composition and metabolic capacity, we hypothesized that human fecal microbiota exhibit pronounced donor-dependent differences in their ability to deconjugate TBBPA-sulfate, resulting in variable regeneration of unconjugated TBBPA and its potential subsequent transformation into Tri-BBPA through microbial debromination. We further hypothesized that the extent of both deconjugation and debromination varies among individuals, potentially influencing TBBPA reactivation and downstream metabolite formation. As a first step to test this hypothesis, we conducted the Pilot-10 Human Donor Microbiome Study, an initial investigation using fecal microbiota from 10 individual human donors (5 male and 5 females) to characterize the capacity and interindividual variability of microbiome-mediated TBBPA-sulfate deconjugation. Specifically, we investigated the extent to which human fecal microbiota metabolizes TBBPA-sulfate and regenerate unconjugated TBBPA, providing mechanistic insight into a potentially overlooked pathway of xenobiotic reactivation and re-toxification. These findings may help establish the biological basis for interindividual differences in TBBPA intestinal exposure and toxicokinetics. TBBPA-glucuronide deconjugation represents an important complementary pathway that warrants future investigation.

## Materials and methods (Note: the experimental design is summarized in Figure 1C)

### Chemicals and Reagents

Individual fecal microbiome samples were obtained from 5 male and 5 female healthy adult human donors through BioIVT (Woodbury, NY). Tetrabromobisphenol A (TBBPA) was purchase from Sigma Alderich (St. Louis, MO) and TBBPA sulfate (S-TBBPA) was from Cayman Chemical Company, Inc. (Ann Arbor, MI). All other chemicals and reagents, unless otherwise noted, were purchased from Sigma Aldrich.

### Human microbiome incubation and chemical exposure

Using a Whitley DG250 Anaerobic Workstation to simulate the gut environment at 37 °C. The human microbiome was prepared by using stool samples to 0.1% L-cysteine, 0.0001% resazurin, and 1.25% glycerol in DPBS to create a slurry. With the slurry, we then used 1g to 1mL of media that contains (DPBS, 0.1% L-cysteine, 0.0001% resazurin). In a 48-well plate format with a total volume of 970uL which contain 950 (100mg/mL) of fecal slurry with 20uL (1mg/mL) of S-TBBPA. To confirm the necessity of microbial enzyme activities in S-TBBPA deconjugation, S-TBBPA or vehicle (DMSO) was added to either unheated microbiome samples, or microbiome samples pre-heated for 10 minutes at 110°C. After six hours of incubation, 1% formic acid was added to each well to quench the reaction, and samples were centrifuged at 4,000g for 10 minutes.

### Human Microbiome Incubation and Chemical Exposure

Human fecal microbiome incubations were performed under anaerobic conditions at 37°C using a Whitley DG250 Anaerobic Workstation to simulate the intestinal environment similar to what we describe previously ^20^. Human stool samples were homogenized in Dulbecco’s phosphate-buffered saline (DPBS) supplemented with 0.1% L-cysteine, 0.0001% resazurin, and 1.25% glycerol to prepare fecal slurries. The slurries were further diluted at a ratio of 1 g fecal material per 1 mL of incubation medium containing DPBS, 0.1% L-cysteine, and 0.0001% resazurin. Incubations were conducted in 48-well plates with a total reaction volume of 970 µL per well, consisting of 950 µL fecal slurry (100 mg/mL) and 20 µL S-TBBPA stock solution (1 mg/mLIn a separate set of experiments, to determine whether microbial enzymatic activity was required for S-TBBPA deconjugation, S-TBBPA or vehicle control (DMSO) was added to either untreated fecal microbiome samples or samples heat-inactivated at 110°C for 10 min. Following incubation for 6 h under anaerobic conditions at 37°C, reactions were quenched by adding formic acid to a final concentration of 1%. Samples were subsequently centrifuged at 4,000 × g for 10 min to remove particulate material.

### Liquid chromatography-mass spectrometry (LC-MS)

Lipids were extracted from microbial samples using a modified methyl tert-butyl ether (MTBE)-based extraction protocol. Briefly, microbial pellets were resuspended and homogenized by sonication in an ice-cold water bath for 30 min to disrupt microbial cells. Subsequently, 50 µL of homogenized sample was transferred into a 1.5 mL solvent-resistant microcentrifuge tube prewashed twice with hexane. Ice-cold phosphate-buffered saline (PBS; 50 µL), methanol (40 µL), and MTBE (200 µL) were sequentially added. Samples were vortexed for 30 s and incubated at −20°C for 30 min, followed by centrifugation at 14,000 rpm for 10 min to separate the organic and aqueous phases. The upper MTBE layer (150 µL) was collected into glass LC-MS vials and evaporated to dryness in a chemical fume hood. Dried lipid extracts were reconstituted in 200 µL of isopropanol:methanol (1:1, v/v), briefly vortexed, and centrifuged at 14,000 rpm for 10 min. The resulting supernatant (150 µL) was transferred into glass vial inserts for LC-MS analysis. Aliquots from individual samples were pooled to generate quality control (QC) samples. S-TBBPA, TBBPA, and tri-BBPA were quantified by LC-MS using an Agilent 1290 UPLC-6495 QQQ-MS system (Santa Clara, CA). Reverse phase chromatography was used with a Waters XSelect HSS T3 column (150 x 2.1 mm, 2.5 µm particle size; Waters Corporation, Milford, MA). Flow through the column was maintained at 0.3 mL/min. The mobile phase Solvent A was comprised of 10 mM ammonium acetate in 60% H_2_O/40% ACN. Solvent B consisted of 10 mM ammonium acetate in 90% IPA/10% ACN. An isocratic elution was used with 50% solvent B for 3 minutes before its percentage was gradually increased to 100% over the next 12 minutes. Following 10 minutes of continued 100% solvent B, at t = 25 minutes, the percent of B was decreased gradually back to 50% to prepare for the next sample injection. The mass spectrometer is equipped with an electrospray ionization (ESI) source. Targeted data acquisition was performed in multiple-reaction-monitoring (MRM) mode. The whole LC-MS system was controlled by Agilent Masshunter Workstation software (Santa Clara, CA). The extracted MRM peaks were integrated using Agilent MassHunter Quantitative Data Analysis (Santa Clara, CA).

### Microbial DNA extraction

DNA Isolation was performed by using an E.Z.N.A. Stool DNA isolation kit (Omega Bio-tek, Norcross, GA). Using 200 mg of the sample in a 2 mL Eppendorf tube with 200 mg of glass beads and SLX-Mlus Buffer then vortex until completely homogenized, followed by adding 30uL DS buffer, 20uL of Proteinase K solution and then incubate for 10mins (70°C;1200 rpm) using an Eppendorf® ThermoMixer (Burlington, MS). Two hundred microliters of SP2 buffer is added and then vortex, chill for 5minutes and centrifuge for 5 minutes (≥ 13,000 x g). Into a new 1.5mL Eppendorf tube 400uL supernatant is added with 200 uL CHTR Reagent. After centrifuging for 2 minutes, equal parts of supernatant, BL buffer, and 100% ethanol into a 2 mL collection tube with HiBind DNA Mini Column. After discarding the filtrate, 500uL of VHB Buffer is added and then wash twice with 700uL of DNA Wash Buffer columns are then transfer into a clean 1.5mL tube with Elution Buffer heated to 65°C. DNA concentration was quantified by a Qubit fluorometer using a dsDNA HS assay kit (Waltham, Massachusetts). DNA was stored at -80°C until further analysis.

### Metagenomic Shotgun Sequencing

Shallow shotgun metagenomic sequencing was performed on human fecal microbiome samples by SailGene Technology USA (Worcester, MA, USA) using 2 × 150 bp paired-end sequencing, targeting approximately 5 million reads per sample. Raw data were trimmed using fastp with a minimum read length of 100 bp, maximum read length of 150 bp, and minimum quality threshold of 20. Taxonomic abundances were estimated using MetaPhlAn 4.1.1 ^21^ using mpa_vJun23_CHOCOPhlAnSGB_202307, and microbial gene-family abundances were quantified using HUMAnN (v3.9) ^22^ with UniRef90 annotations. Differential abundance analysis was performed using EdgeR (FDR-BH < 0.05).

### Hierarchical clustering

Heatmaps and hierarchical clustering analyses were performed in Python using SciPy, pandas, Matplotlib, and Seaborn. For LC-MS metabolite profiling, two-way hierarchical clustering was performed using Euclidean distance and average linkage. The TBBPA/S-TBBPA ratio (%) was calculated as (TBBPA concentration/S-TBBPA concentration) × 100 and displayed separately with an independent color scale. For microbial functional profiling, gene families annotated as arylsulfatases, other sulfatases, or sulfatase-maturation/activation enzymes were identified from HUMAnN v3.9 results using UniRef90 annotations. Gene families with mean abundance greater than 1 count per million (CPM) across donors were retained. For each functional category, gene-family abundances were standardized using row-wise Z-scores, followed by two-way hierarchical clustering of gene families and donors using Euclidean distance and average linkage. Heatmaps were visualized using a blue-white-red color scale, with red indicating higher and blue indicating lower relative abundance.

### Spearman’s correlation analyses

Correlations between microbial gene abundance and TBBPA-sulfate deconjugation activity were assessed using Spearman’s rank correlation. Analyses were restricted to nine donors with matched LC-MS and metagenomic data (M1-M5 and F1-F4). For category-level analyses, DNA abundances of all annotated gene families within each functional category were summed for each donor to obtain category-specific cumulative abundance. These values were correlated with the corresponding TBBPA/S-TBBPA ratios (%). Individual UniRef90 gene-family abundances, including taxonomically stratified annotations, were also correlated with the metabolite ratios to identify candidate microbial enzymes associated with deconjugation activity. Two-sided permutation tests accounting for tied abundance values were used to calculate exact Spearman *p* values. Scatterplots were generated with individual donors represented as points with linear regression lines,

## Results

### Human fecal microbiota mediate TBBPA-sulfate deconjugation without detectable debromination

As shown **in Figure 2A**, anaerobic incubation of TBBPA-sulfate with human fecal microbiota at 37 °C (unheated), which simulated the human gut environment, resulted in a significant decrease in TBBPA-sulfate concentrations, accompanied by the accumulation of unconjugated TBBPA, indicating active microbial deconjugation. In contrast, fecal microbiota subjected to heat inactivation at 95°C (heated) exhibited negligible TBBPA formation, with most TBBPA-sulfate remaining intact. These findings indicate that TBBPA-sulfate deconjugation is dependent on heat-sensitive microbial enzyme activity rather than spontaneous chemical hydrolysis. Notably, Tri-BBPA remained undetectable or negligible in both heated and unheated samples, suggesting that microbial debromination did not occur to a measurable extent under the experimental conditions in the human fecal microbiota. As shown in **Figure 2B**, sex-stratified analysis of unheated fecal incubations demonstrated TBBPA-sulfate depletion and concomitant TBBPA formation in samples from both male and female donors. Although the concentrations of residual TBBPA-sulfate and regenerated TBBPA varied among individual donors, no statistically significant sex-dependent differences were observed. Tri-BBPA remained undetectable or negligible in both groups, further indicating the absence of measurable debromination. Collectively, these results demonstrate that human fecal microbiota has heat-sensitive TBBPA-sulfate deconjugation enzymatic activity, with interindividual variability but no detectable sex-associated differences or appreciable downstream debromination under the conditions and sample size examined.

**Figure 2.**
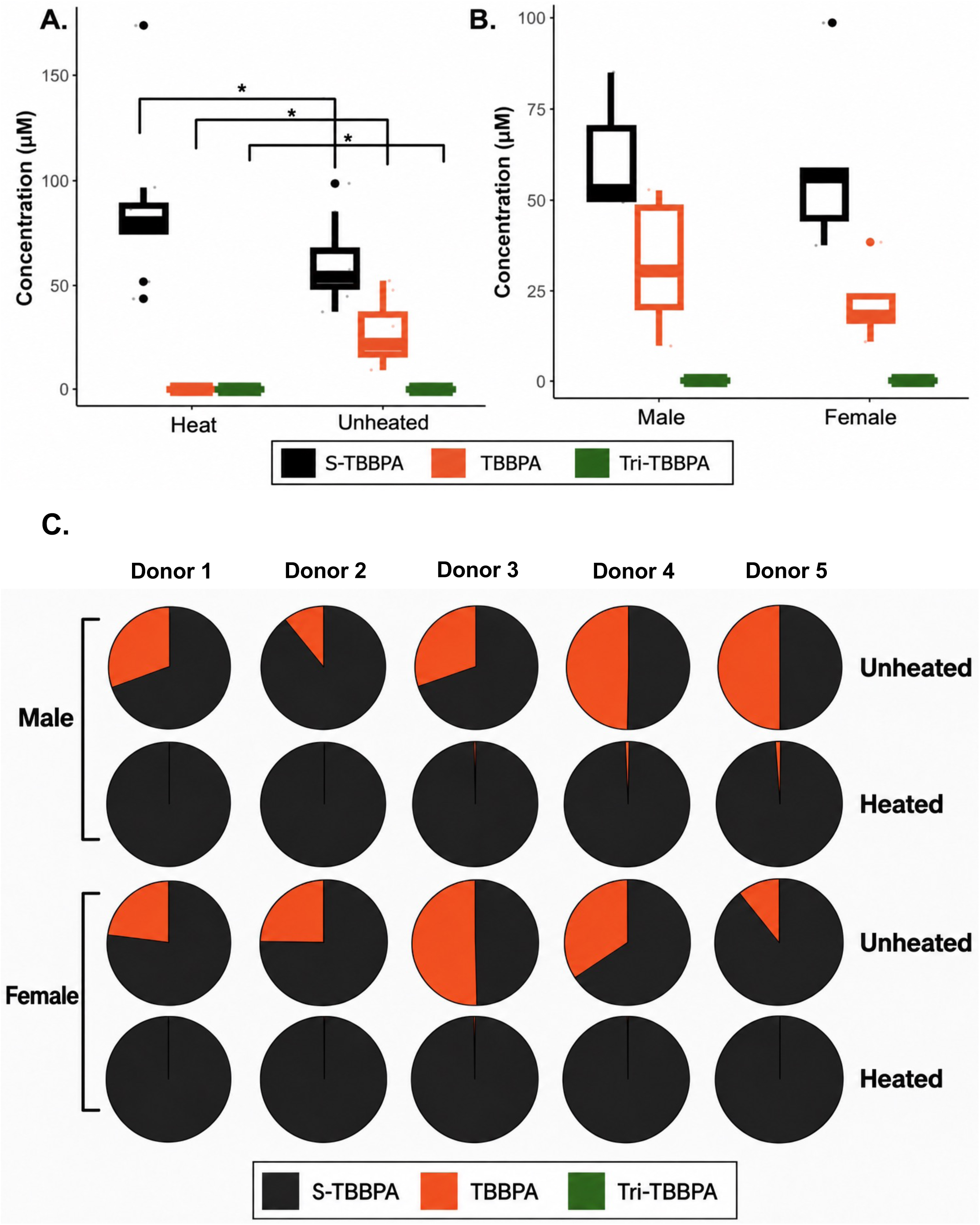
Heat-sensitive microbial deconjugation of TBBPA-sulfate and interindividual variability in metabolite profiles. **A.** LC-MS quantification of TBBPA-sulfate (S-TBBPA), unconjugated TBBPA, and tribromobisphenol A (Tri-BBPA, a potential downstream metabolite of TBBPA) in heat-treated and unheated human fecal microbiome cultures following incubation with TBBPA-sulfate. Unheated cultures exhibited substantial conversion of TBBPA-sulfate to TBBPA, whereas TBBPA formation was markedly reduced or nearly abolished following heat inactivation, supporting the involvement of heat-sensitive microbial enzymatic processes. Tri-BBPA was undetectable or negligible under the experimental conditions, suggesting minimal downstream debromination. **B.** Comparison of TBBPA-sulfate, TBBPA, and Tri-BBPA concentrations between male and female donor microbiome cultures. No significant sex-associated differences in metabolite profiles were observed in this pilot cohort. **C.** Donor-specific distributions of TBBPA-sulfate, TBBPA, and Tri-BBPA in unheated and heat-treated fecal microbiome cultures from five male and five female donors. Pie charts illustrate the relative proportions of the three measured metabolites within each sample. Substantial interindividual variability in TBBPA-sulfate conversion was observed among unheated cultures, whereas heat-treated controls remained predominantly composed of TBBPA-sulfate, with minimal TBBPA formation. Black, orange, and green represent TBBPA-sulfate, TBBPA, and Tri-BBPA, respectively.

As shown in **Figure 2C**, to further characterize the TBBPA-sulfate deconjugation at the individual donor level, pie charts were plotted among the 10 human fecal microbiota samples (five male [M1-M5] and five female [F1-F5] donors). In unheated incubations, the proportion of regenerated TBBPA varied markedly across donors, ranging from approximately 10% to 50% of the measured compounds, indicating heterogeneous microbial deconjugation capacities. In contrast, heat-inactivated (95°C) samples exhibited nearly exclusive recovery of TBBPA-sulfate with negligible TBBPA formation, supporting the involvement of heat-sensitive microbial activity. Notably, substantial differences in deconjugation were observed within both male and female donor groups, consistent with donor-dependent rather than sex-specific variation. Tri-BBPA was not detected at appreciable levels in any individual samples. Collectively, these findings highlight substantial interindividual heterogeneity in the capacity of human fecal microbiota to regenerate unconjugated TBBPA from TBBPA-sulfate.

As shown in **Figure 3**, two-way hierarchical clustering dendrogram demonstrated distinct putative microbial metabolizer phenotypes. Unheated samples from high and intermediate metabolizers generally clustered separately from heat-inactivated controls, although some low-metabolizer samples clustered with the heat-inactivated group, consistent with active microbial deconjugation in unheated samples and minimal TBBPA formation following heat inactivation at 95°C. Based on the relative abundance of regenerated TBBPA and residual TBBPA-sulfate (S-TBBPA), together with the TBBPA/S-TBBPA ratio (%) at the experimental endpoint, donors were categorized into three putative metabolizer groups: high metabolizers (F3, M4, and M5), intermediate metabolizers (M1, F1, F2, F4, and M3), and low metabolizers (F5 and M2). High metabolizers exhibited the highest TBBPA/S-TBBPA ratios (97.1 ∼ 100%), indicating substantial TBBPA accumulation relative to residual S-TBBPA, whereas low metabolizers showed the lowest ratios (11.1 ∼ 11.3%), reflecting limited TBBPA formation. Intermediate metabolizers displayed moderate conversion profiles, with ratios ranging from 28.6% to 52.1%. Notably, Tri-BBPA remained at comparatively low concentrations across samples, indicating that it is not a significant downstream metabolite of the human microbiome. All three groups included donors of both sexes, indicating that the observed metabolic heterogeneity was not exclusively associated with sex. Collectively, these findings highlighted substantial donor-dependent variation in the capacity of human fecal microbiota to deconjugate TBBPA-sulfate, potentially contributing to interindividual differences in intestinal TBBPA regeneration and exposure.

**Figure 3.**
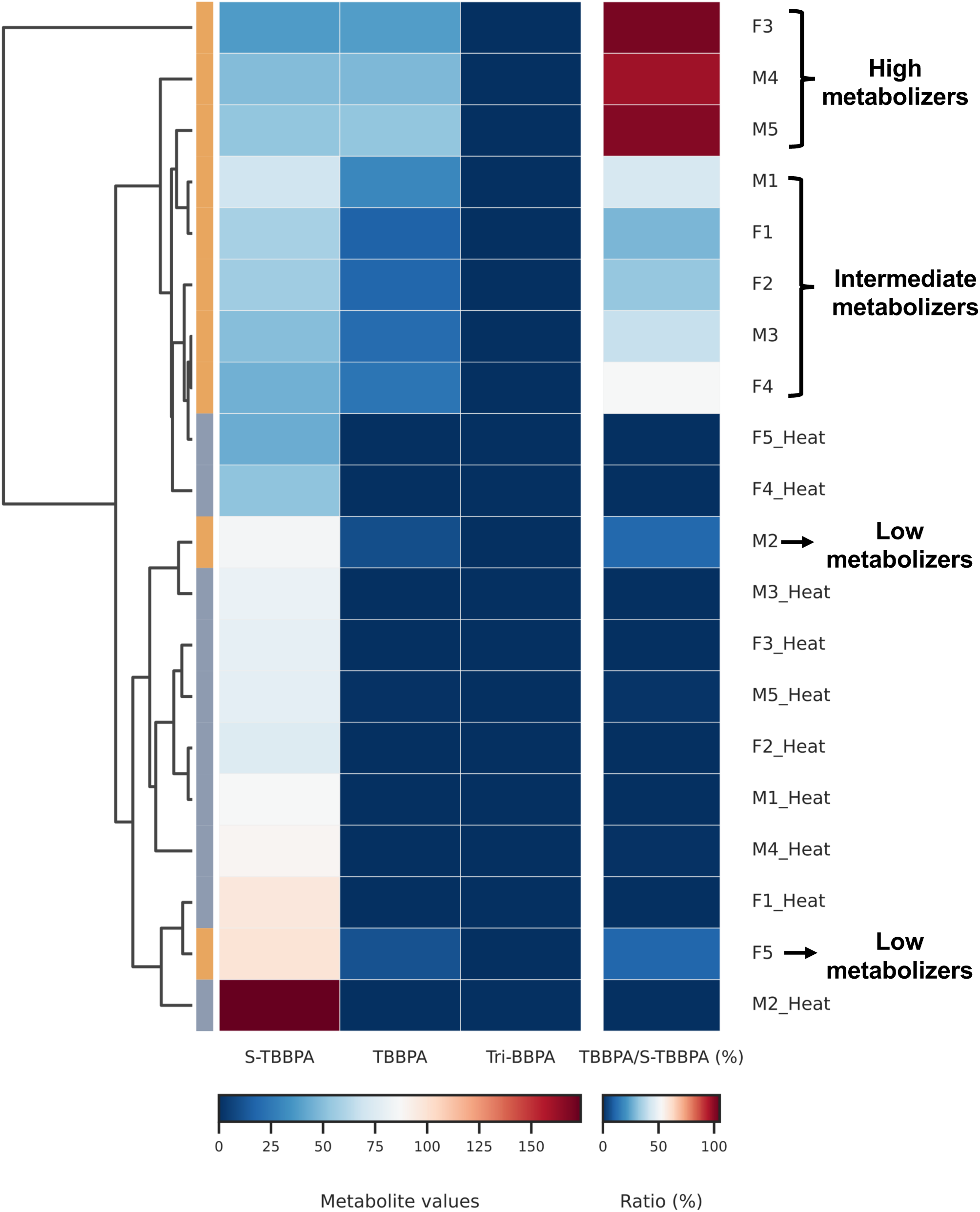
Unsupervised hierarchical clustering illustrating donor-specific LC-MS metabolite profiles of TBBPA-sulfate and TBBPA in unheated human fecal microbiome cultures and their corresponding heat-treated controls. Hierarchical clustering heatmap of LC-MS-measured S-TBBPA, TBBPA, and Tri-BBPA concentrations in unheated and heat-inactivated human fecal microbiome cultures following 6-hour incubation. Samples were clustered using Euclidean distance and average linkage, with metabolite concentrations displayed using a blue–white–red color scale. The TBBPA/S-TBBPA ratio (%) was displayed as a separate column with an independent color scale to illustrate donor-specific deconjugation activity. Donors exhibited substantial interindividual differences in metabolite profiles, with high, intermediate, and low TBBPA-producing phenotypes, whereas heat-inactivated cultures showed minimal TBBPA formation. These findings highlight the marked heterogeneity and heat-sensitive nature of microbiome-mediated S-TBBPA deconjugation.

### Donor-specific microbial arylsulfatase gene abundance and correlation with TBBPA-sulfate deconjugation activity

The substantial interindividual variability in TBBPA-sulfate deconjugation observed by LC-MS led us to investigate the underlying microbial genetic determinants of this metabolic activity. Given that sulfatases catalyze the hydrolysis of sulfate ester bonds, we first examined whether differences in microbial arylsulfatase gene abundance could explain the donor-dependent variation in TBBPA-sulfate deconjugation. To address this question, we performed MGS of fecal microbiome samples from nine donors (to note, female donor 5 was not subjected to MGS due to low DNA yield), followed by microbial gene-family profiling using HUMAnN v3.9 with UniRef90 annotations, as described in Materials and Methods. Arylsulfatase gene families with a mean abundance greater than 1 count per million (CPM) across donors were selected for hierarchical clustering and association analyses. As shown in **Figure 4A**, two-way hierarchical clustering revealed substantial donor-dependent heterogeneity in microbial arylsulfatase gene abundance, with distinct patterns associated with different bacterial taxa. Arylsulfatase genes associated with *Eubacterium hallii* exhibited relatively high abundance in female donor # 2 (F2), male donor # 4 (M4), and M5, whereas genes associated with *Roseburia inulinivorans*, *Coprococcus comes*, and *Phascolarctobacterium faecium* were predominantly enriched in M1, M4, and M5. In contrast, multiple *Escherichia coli*-associated arylsulfatase genes showed pronounced enrichment in M2, with additional enrichment in F2. Arylsulfatase genes associated with *Bacteroides vulgatus* also exhibited relatively high abundance in F2, whereas those associated with *Blautia obeum*, *Bifidobacterium longum*, and *Anaerostipes hadrus* were enriched primarily in F1 and M3. Furthermore, *Eggerthella lenta*-associated arylsulfatase genes exhibited increased relative abundance in F3 and F4. These patterns demonstrate substantial interindividual variation in the taxonomic distribution and relative abundance of microbial arylsulfatase genes. As shown in **Figure 4B**. Spearman correlation analysis identified five microbial arylsulfatase gene-family annotations whose DNA abundances were positively associated with the TBBPA/S-TBBPA LC-MS metabolite ratio (%) across nine human fecal microbiome donors. Four arylsulfatase genes assigned to *Firmicutes bacterium CAG 114* (UniRef90_R5HNY1, R5HAJ4, R5HNT9, and R5HPX2) exhibited significant positive correlations (Spearman’s ρ = 0.745 ∼ 0.795, P = 0.0144 ∼ 0.0268). An additional arylsulfatase gene from *Bacteroides thetaiotaomicron* (UniRef90_A0A139L2J0) also showed a positive association (ρ = 0.686, P = 0.0475). These findings identified candidate microbial arylsulfatases potentially associated with interindividual variability in TBBPA-sulfate deconjugation.

**Figure 4.**
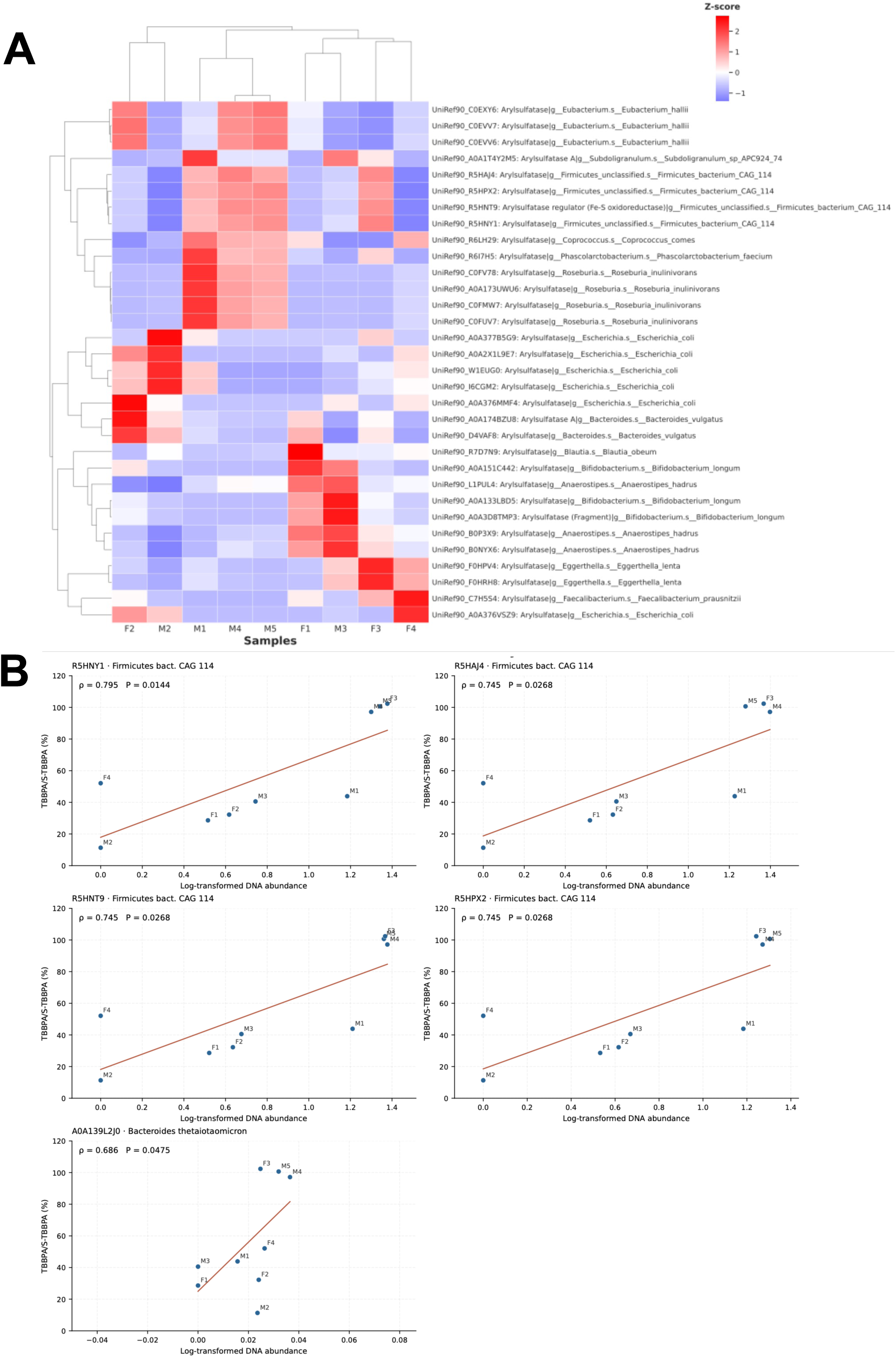
Microbial arylsulfatase gene abundance and its association with TBBPA-sulfate deconjugation activity. **A.** Two-way hierarchical clustering heatmap of microbial arylsulfatase gene abundances across nine human fecal microbiome samples. MGS-derived microbial gene-family abundances were quantified using HUMAnN v3.9 with UniRef90 annotations. Gene families annotated as arylsulfatases with mean abundance greater than 1 CPM across donors were retained, yielding 32 taxonomically stratified gene-family entries. Row-wise Z-scores were calculated from the original, non-log-transformed abundance values. Hierarchical clustering of both gene families and donors was performed using Euclidean distance and average linkage. Red indicates higher and blue indicates lower relative gene abundance across donors. **B.** Individual Spearman correlations between microbial arylsulfatase gene-family DNA abundance and LC-MS-measured TBBPA/S-TBBPA ratio (%) across nine donors. Five taxonomically stratified arylsulfatase annotations exhibited nominally significant positive associations with deconjugation activity (Spearman’s ρ = 0.686 - 0.795; *p* < 0.05). Each point represents an individual donor, and red lines indicate fitted linear trends. These associations identify candidate microbial arylsulfatases potentially contributing to donor-dependent TBBPA-sulfate deconjugation.

### Other microbial sulfatase gene abundance and correlation with TBBPA-sulfate deconjugation activity

We next expanded our analysis to other microbial sulfatases that may contribute to the hydrolysis of sulfate-conjugated compounds. These included genes annotated as choline sulfatases, putative sulfatases, sulfatase family proteins, and other uncharacterized sulfatases. Using HUMAnN v3.9 with UniRef90 annotations, we identified 21 taxonomically stratified sulfatase gene-family entries with a mean abundance greater than 1 CPM across donors. As shown in **Figure 5A**, two-way hierarchical clustering revealed substantial donor-dependent heterogeneity in the abundance of other microbial sulfatase genes, with a pronounced contribution from *Escherichia coli*-associated gene families. Among the 21 identified entries, 18 were associated with *E. coli*, including genes annotated as choline sulfatases, putative sulfatases, the uncharacterized sulfatase YidJ, and the putative alkyl/aryl-sulfatase YjcS. These *E. coli*-associated genes were particularly enriched in M2, with additional enrichment in F3, F2, and M1, whereas their abundances were markedly lower or undetectable in F1, M4, and M5. In contrast, sulfatase genes associated with *Roseburia inulinivorans* were relatively enriched in M1, M4, and M5, while those associated with *Ruminococcus torques* and *Eubacterium rectale* exhibited higher abundances in F3 and F4. These distinct patterns demonstrate considerable interindividual variation in the taxonomic distribution and abundance of microbial sulfatase genes beyond those annotated specifically as arylsulfatases. As shown in **Figure 5B**, Spearman correlation analysis identified two taxonomically stratified annotations of the microbial sulfatase gene UniRef90_A0A2U2EKR0 whose DNA abundances were positively associated with the TBBPA/S-TBBPA LC-MS metabolite ratio (%) across nine human fecal microbiome donors. The gene annotations assigned to *Ruminococcus torques* and *Eubacterium rectale* exhibited significant positive correlations (Spearman’s ρ = 0.850, *p* = 0.0061 and ρ = 0.767, *p* = 0.0214, respectively). These findings identified additional candidate microbial sulfatase genes associated with interindividual variability in TBBPA-sulfate deconjugation.

**Figure 5.**
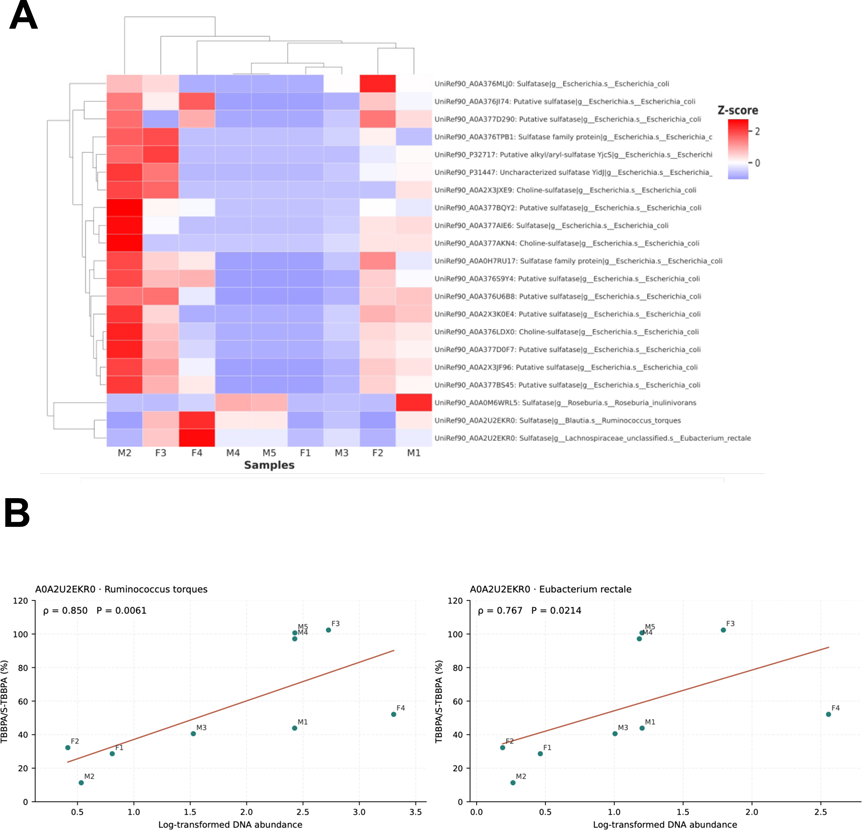
Other microbial sulfatase gene abundance and its association with TBBPA-sulfate deconjugation activity. **A.** Two-way hierarchical clustering heatmap of other microbial sulfatase gene abundances across nine human fecal microbiome samples. Gene-family abundances were quantified using HUMAnN v3.9 with UniRef90 annotations. Sulfatase gene families other than those classified as arylsulfatases or sulfatase-maturation enzymes were extracted, and entries with mean abundance greater than 1 CPM across donors were retained, yielding 21 taxonomically stratified gene-family entries. Row-wise Z-scores were calculated from original, non-log-transformed abundance values, followed by hierarchical clustering of both gene families and donors using Euclidean distance and average linkage. Red indicates higher and blue indicates lower relative gene abundance. **B.** Individual Spearman correlations between other microbial sulfatase gene-family DNA abundance and the LC-MS-measured TBBPA/S-TBBPA ratio (%) across nine donors. Two taxonomically stratified annotations of UniRef90_A0A2U2EKR0, assigned to *Ruminococcus torques* and *Eubacterium rectale*, exhibited nominally significant positive associations with deconjugation activity (ρ = 0.850, *p* = 0.0061 and ρ = 0.767, *p* = 0.0214, respectively). Each point represents an individual donor, and red lines indicate fitted linear trends. These findings identify additional microbial sulfatase candidates associated with TBBPA-sulfate deconjugation.

### Microbial sulfatase-maturation enzyme DNA abundance and correlation with TBBPA-sulfate deconjugation activity

We next investigated whether microbial sulfatase-maturation enzymes could explain the observed donor-dependent differences in metabolic capacity. To note, many bacterial sulfatases require post-translational conversion of a conserved cysteine or serine residue into catalytically essential formylglycine to become functionally active. Under anaerobic conditions, this modification can be catalyzed by anaerobic sulfatase-maturation enzymes (anSMEs) ^23–25^. We therefore examined whether the abundance of genes encoding sulfatase-maturation or activation enzymes was associated with LC-MS-measured TBBPA-sulfate deconjugation activity. Using HUMAnN v3.9 with UniRef90 annotations, we identified 33 taxonomically stratified sulfatase-maturation or activation enzyme gene-family entries with a mean abundance greater than 1 CPM across donors. As shown in **Figure 6A**, two-way hierarchical clustering revealed distinct donor-specific patterns of microbial sulfatase-maturation enzyme gene abundance. Genes encoding anaerobic sulfatase-maturating enzyme homologs, particularly YdeM-associated proteins, were distributed across multiple bacterial taxa, including *Blautia wexlerae*, *Blautia obeum*, *Anaerostipes hadrus*, *Roseburia inulinivorans*, *Roseburia hominis*, *Coprococcus comes*, *Ruminococcus torques*, and *Eubacterium rectale*. Notably, M4 and M5 exhibited relatively high abundances of several YdeM-associated gene families originating from *Blautia*, *Anaerostipes*, and *Coprococcus*, whereas F1 showed enrichment of maturation-enzyme genes associated with *Roseburia hominis*, *Ruminococcus torques*, and *Fusicatenibacter saccharivorans*. In contrast, F4 exhibited pronounced enrichment of maturation-enzyme genes associated with *Eubacterium rectale* and *Ruminococcus torques*. Several *Escherichia coli*-associated genes encoding anaerobic sulfatase-maturation enzymes, putative arylsulfatase-activating proteins, and AtsB-domain proteins were relatively enriched in M2 and F3. These findings demonstrate substantial interindividual heterogeneity in the taxonomic origins and relative abundance of microbial sulfatase-maturation genes.

**Figure 6.**
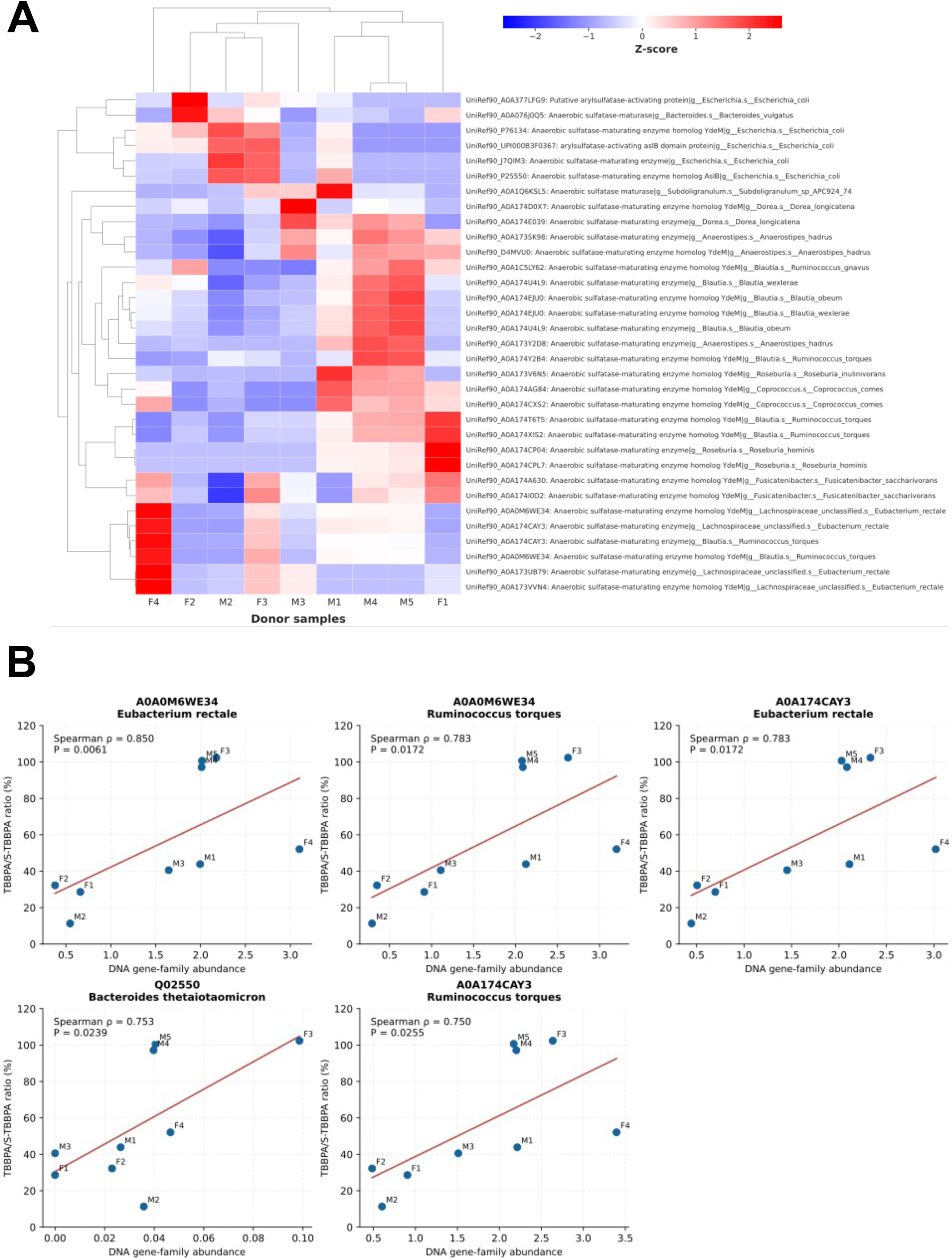
Microbial sulfatase-maturation enzyme gene abundance is positively associated with TBBPA-sulfate deconjugation activity. **A.** Two-way hierarchical clustering heatmap of microbial sulfatase-maturation or activation enzyme (anSME) gene abundances across nine human fecal microbiome samples. Gene-family abundances were quantified using HUMAnN v3.9 with UniRef90 annotations. Entries annotated as sulfatase-maturation or activation enzymes with mean abundance greater than 1 CPM across donors were retained, yielding 33 taxonomically stratified gene-family entries. Row-wise Z-scores were calculated from original, non-log-transformed abundance values, followed by hierarchical clustering of both gene families and donors using Euclidean distance and average linkage. Red indicates higher and blue indicates lower relative gene abundance. **B.** Individual Spearman correlations between sulfatase-maturation enzyme gene-family DNA abundance and the LC-MS-measured TBBPA/S-TBBPA ratio (%) across nine donors. Five taxonomically stratified gene-family annotations exhibited nominally significant positive correlations (ρ = 0.750 - 0.850; *p* = 0.0061 - 0.0255), including UniRef90_A0A0M6WE34 and UniRef90_A0A174CAY3 assigned to *Eubacterium rectale* and *Ruminococcus torques*, and UniRef90_Q02550 assigned to *Bacteroides thetaiotaomicron*. Each point represents an individual donor, and red lines indicate fitted linear trends. These associations support sulfatase maturation capacity as a candidate determinant of interindividual variability in TBBPA-sulfate deconjugation.

As shown in **Figure 6B**, Spearman correlation analysis identified five taxonomically stratified microbial sulfatase-maturation enzyme gene-family annotations whose DNA abundances were positively assoc4iated with the TBBPA/S-TBBPA ratio (%) across nine human fecal microbiome donors. Two annotations of UniRef90_A0A0M6WE34, assigned to *Eubacterium rectale*and *Ruminococcus torques*, exhibited significant positive correlations (Spearman’s ρ = 0.850, *p* = 0.0061 and ρ = 0.783, *p* = 0.0172, respectively). Similarly, UniRef90_A0A174CAY3 annotations from *E. rectale* and *R. torques* were positively correlated with the metabolite ratio (ρ = 0.783, *p* = 0.0172 and ρ = 0.750, *p* = 0.0255, respectively). An additional sulfatase-maturation enzyme annotation, UniRef90_Q02550 from *Bacteroides thetaiotaomicron*, also showed a significant positive association (ρ = 0.753, *p* = 0.0239). Collectively, these findings identified candidate microbial sulfatase-maturation enzymes associated with interindividual variability in TBBPA-sulfate deconjugation activity.

To determine whether the collective abundance of microbial sulfatase-related genes better reflects TBBPA-sulfate deconjugation activity than individual gene abundances, we next performed category-level correlation analyses. Because multiple microbial sulfatases may contribute to xenobiotic deconjugation through functional redundancy, we reasoned that aggregating gene abundances within each functional category could provide a more comprehensive measure of the microbial community’s potential for desulfation. Accordingly, we evaluated the cumulative DNA abundance of three functional categories: arylsulfatases, other sulfatases, and sulfatase-maturation/activation enzymes, in relation to the TBBPA/S-TBBPA ratio across nine donors with matched metagenomic and LC-MS data. As shown in **Figure 7A**, sulfatase-maturation/activation enzyme abundance exhibited a significant positive association with the TBBPA/S-TBBPA ratio (Spearman’s ρ = 0.717, *p* = 0.037). In contrast, neither arylsulfatase abundance (ρ = −0.167, *p* = 0.678) nor other sulfatase abundance (ρ = −0.100, *p* = 0.810) showed a significant correlation. These findings suggest that the collective abundance of sulfatase-maturation enzymes may be a more informative indicator of microbial TBBPA-sulfate deconjugation activity than sulfatase gene abundance alone, highlighting the potential importance of enzyme maturation and activation in determining microbial desulfation capacity. Although this association does not necessarily mean causation, our findings support further investigation of microbial sulfatase maturation as a potential functional determinant of TBBPA-sulfate deconjugation and interindividual variability in intestinal TBBPA regeneration.

**Figure 7.**
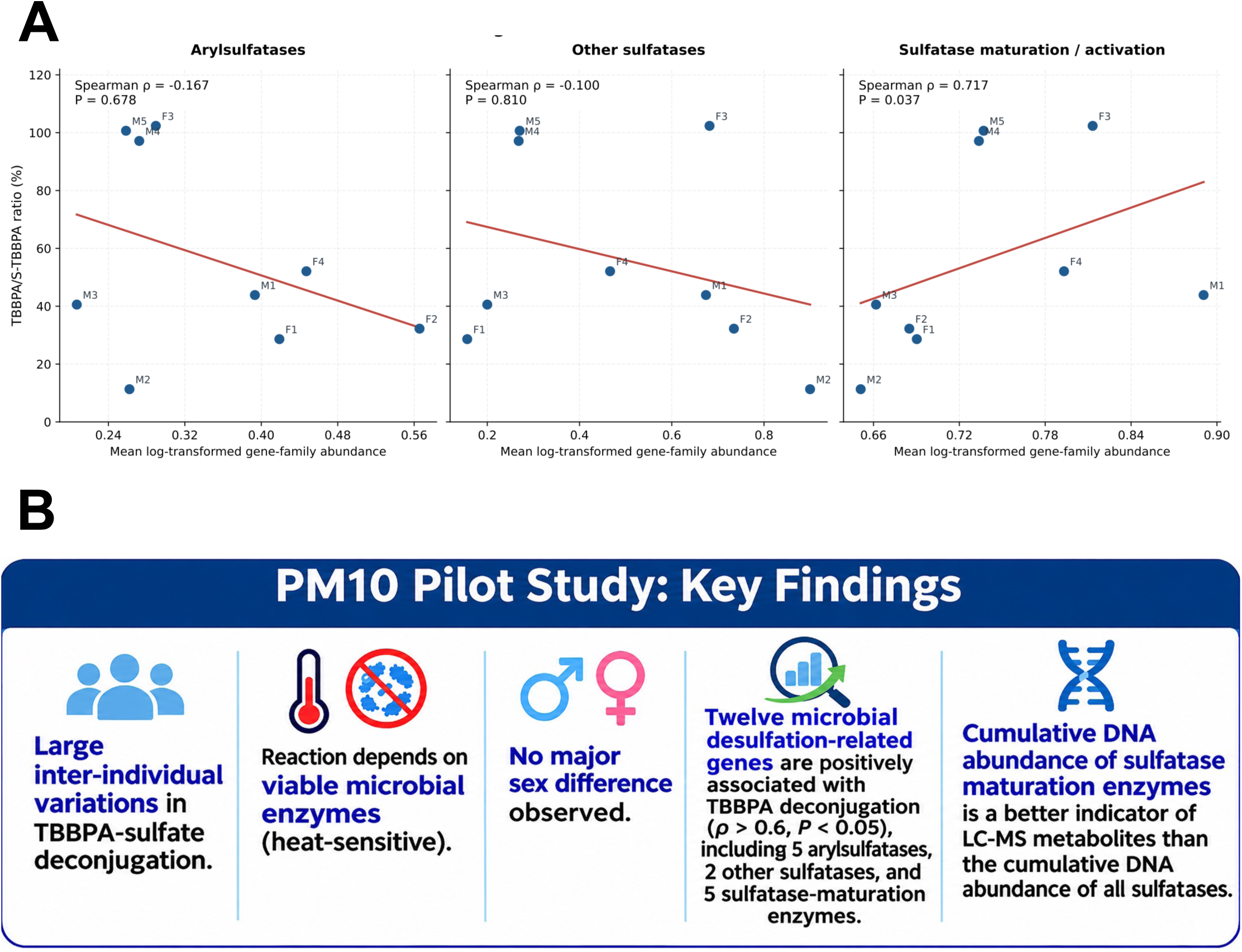
Integrated associations between cumulative DNA abundance of microbial sulfatase-related gene families and and TBBPA-sulfate deconjugation activity, and graphical summary of the PM10 pilot study. **A.** For each donor, DNA abundances of all annotated gene families within each functional category (arylsulfatases, other sulfatases, or sulfatase-maturation/activation enzymes) were summed to calculate the cumulative DNA abundance for that category. Spearman’s rank correlations were then performed between category-specific cumulative DNA abundance and the LC-MS-measured TBBPA/S-TBBPA ratio (%) across nine human fecal microbiome donors. The cumulative DNA abundance of the sulfatase-maturation/activation enzyme family exhibited a positive association with deconjugation activity (Spearman’s ρ = 0.717, *p* = 0.037), whereas arylsulfatases (ρ = −0.167, *p* = 0.678) and other sulfatases (ρ = −0.100, *p* = 0.810) DNA abundances showed no significant associations. Each point represents an individual donor, and fitted lines illustrate overall trends. **B.** Graphical summary of five major findings from the Pilot-10 Human Microbiome Donor Study (PM10): (1) substantial interindividual variability in S-TBBPA deconjugation; (2) marked suppression of TBBPA regeneration following heat inactivation, supporting heat-sensitive microbial enzymatic activity; (3) no major sex-associated differences in this pilot cohort; (4) identification of 12 taxonomically stratified microbial desulfation-related gene-family annotations positively associated with TBBPA/S-TBBPA ratios (ρ > 0.6, nominal P < 0.05), including five arylsulfatases, two other sulfatases, and five sulfatase-maturation enzyme annotations; and (5) a stronger association of sulfatase-maturation enzyme DNA abundance with TBBPA-sulfate deconjugation than overall sulfatase gene abundance. Collectively, these findings suggest that microbial sulfatase-maturation capacity may be a more informative indicator of TBBPA reactivation potential than sulfatase gene abundance alone. These exploratory associations require validation in larger human cohorts and functional experiments. Disclaimer: The experimental workflow illustration in Fig. 7B was created using generative AI and subsequently reviewed and edited by the authors to ensure scientific accuracy. Generative AI was used solely for graphical illustration and was not involved in scientific data generation, analysis, or interpretation.

## Discussion

In conclusion, as shown in **Figure 7B**, our PM10 pilot study delivered 5 major findings: (1) we observed substantial interindividual variability in human microbiome-mediated TBBPA-sulfate deconjugation; (2) we discovered the dependence of TBBPA regeneration on heat-sensitive microbial enzymatic activity; (3) we found no major sex-associated differences in TBBPA-sulfate deconjugation between male and female human donors at the current sample size; (4) we identified 12 microbial desulfation-related gene-family annotations positively associated with the TBBPA/S-TBBPA ratio (ρ > 0.6, nominal *p* < 0.05), including five arylsulfatases, two other sulfatases, and five sulfatase-maturation enzymes; and (5) we discovered a stronger association between sulfatase-maturation enzyme abundance and TBBPA-sulfate deconjugation activity than between overall sulfatase abundance and deconjugation activity. Collectively, these findings highlight sulfatase maturation as a potentially important determinant of interindividual variability in microbial TBBPA reactivation. Although the small sample size and lack of direct enzyme perturbation limit mechanistic conclusions, our PM10 pilot study established a foundation for future studies in identifying the responsible microbial enzymes and developing precision microbiome-directed strategies to limit TBBPA reactivation and potentially reduce internal exposure and associated health risks.

Our findings extend our previous investigation of gut microbiome-mediated deconjugation of polychlorinated biphenyl sulfates (PCB-sulfates), demonstrating that microbial desulfation is relevant to multiple structurally distinct classes of environmental contaminants. In our earlier study ^20^, anaerobically cultured mouse and two human fecal microbiota samples hydrolyzed four PCB-sulfate congeners (PCB3, PCB11, PCB25, and PCB52 sulfates) to their corresponding hydroxylated PCBs (OH-PCBs). Notably, higher-chlorinated PCB-sulfates exhibited greater microbial transformation, demonstrating substrate-dependent differences in deconjugation. We also observed greater PCB-sulfate hydrolysis in male than female mouse microbiota, whereas two human microbial communities exhibited distinct metabolic capacities. Building upon these findings, the present Pilot-10 Human Donor Study expanded the investigation to TBBPA-monosulfate, a structurally distinct brominated flame-retardant metabolite, and examined interindividual variability across a larger number of human donors. Consistent with our previous observations, human fecal microbiota actively deconjugated TBBPA-sulfate, regenerating the unconjugated parent compound. Importantly, we identified substantial donor-dependent differences, with putative fast, intermediate, and slow metabolizer phenotypes, but no statistically significant sex-associated differences. Together, these two studies suggest that microbial desulfation may represent a broader mechanism of xenobiotic reactivation that extends beyond individual chemical classes. The distinct substrate structures and donor-dependent metabolic profiles further highlighted the importance of investigating microbial sulfatase specificity and functional diversity to understand interindividual differences in environmental toxicant metabolism and potential susceptibility.

The present study focused exclusively on TBBPA-monosulfate (S-TBBPA), whereas it is important to note that TBBPA undergoes multiple phase II conjugation reactions that generate a diverse range of metabolites. Previous human biomonitoring studies have identified TBBPA-monoglucuronide, TBBPA-diglucuronide, TBBPA-monosulfate, and TBBPA-disulfate ^9, 12^. In addition, animal studies have identified mixed glucuronide-sulfate conjugates, indicating that TBBPA metabolism involves both individual and sequential conjugation pathways ^9, 11^. These metabolites may differ in their biliary transport, intestinal availability, susceptibility to microbial deconjugation, and potential for enterohepatic recycling. Importantly, the ability of human gut microbiota to hydrolyze these different TBBPA conjugates remains incompletely understood. Our findings as an initial step demonstrated that human fecal microbiota can regenerate unconjugated TBBPA from TBBPA-monosulfate, with substantial donor-dependent variability. However, these results cannot be generalized to TBBPA-disulfate, glucuronide conjugates, or mixed sulfate-glucuronide metabolites, which may require different microbial enzymes or sequential deconjugation reactions. Future studies should systematically compare the microbial transformation of these conjugates and investigate their associated sulfatases, β-glucuronidases, and other relevant enzymes. Such investigations will provide a more comprehensive understanding of how the human gut microbiome influences TBBPA metabolism, enterohepatic cycling, and interindividual variability in internal exposure.

An important limitation of the present study is the relatively small sample size of 10 human fecal donors (five males and five females), which limits the statistical power to evaluate sex-associated differences and the generalizability of the findings to broader human populations. As an initial proof-of-concept investigation, the Pilot-10 Human Donor Study was designed to establish whether human fecal microbiota can deconjugate TBBPA-monosulfate and to explore potential interindividual variability in this metabolic activity. Despite the limited sample size, our findings demonstrated consistent regeneration of unconjugated TBBPA across all 10 donors, negligible deconjugation following heat inactivation, and substantial donor-dependent differences in deconjugation capacity. These observations provide initial experimental evidence supporting human gut microbiome-mediated TBBPA-sulfate reactivation and reveal distinct putative fast, intermediate, and slow metabolizer phenotypes. Nevertheless, these preliminary classifications require validation in larger, more diverse populations and through time-resolved kinetic measurements. Our future studies integrating metagenomic shotgun sequencing, microbial functional analyses, and targeted enzyme validation are being performed to identify the microbial taxa and sulfatases responsible for TBBPA-sulfate deconjugation. Expansion to a larger human donor cohort will further determine the reproducibility of these metabolic phenotypes and their associations with demographic and microbiome characteristics. Despite its exploratory nature, this pioneering study establishes a foundation for understanding interindividual variability in microbiome-mediated TBBPA reactivation and highlights a previously underexplored mechanism through which the gut microbiome may influence environmental toxicant exposure and susceptibility.

Although our findings demonstrated heat-sensitive microbial deconjugation of TBBPA-sulfate, the specific enzymes responsible remain unknown. Microbial arylsulfatases represent plausible candidates because of their capacity to hydrolyze sulfate esters, while anaerobic sulfatase-maturating enzymes (anSMEs) are required for the activation of certain bacterial sulfatases. Previous studies demonstrated that genetic deletion of anSME in the prominent human gut symbiont *Bacteroides thetaiotaomicron* abolished sulfatase activity, whereas deletion of the individual sulfatase BT1636 impaired the utilization of sulfated mucin glycans ^26, 27^. These findings provide a strong rationale for investigating sulfatase-dependent mechanisms underlying TBBPA-sulfate deconjugation. Future studies integrating recombinant enzyme characterization and targeted loss-of-function approaches will be essential to establish their catalytic activity and necessity. Antisense oligonucleotide-based inhibition, particularly peptide nucleic acids, may offer an additional strategy for suppressing selected bacterial genes, although delivery efficiency and target specificity require careful validation ^28, 29^. Chemical inhibition may also provide complementary evidence; however, recent findings demonstrate that certain arylsulfamate compounds affect gut bacterial growth through sulfatase-independent mechanisms, emphasizing the importance of specificity controls ^30^. Collectively, these mechanistic approaches in future studies will help identify the microbial enzymes responsible for TBBPA-sulfate deconjugation and establish whether variation in their abundance or activity contributes to donor-dependent differences in TBBPA regeneration.

Our MGS analyses revealed a significant positive association between cumulative microbial sulfatase-maturation enzyme (anSME) gene family abundance and LC-MS-measured TBBPA-sulfate deconjugation activity (Spearman’s ρ = 0.717, *p* = 0.037;), whereas arylsulfatase (ρ = -0.167, *p* = 0.678) and other sulfatase gene abundances (ρ = -0.100, *p* = 0.810) showed no significant associations. However, at the individual gene-family level, we further identified 12 taxonomically stratified microbial desulfation-related annotations positively correlated with TBBPA-sulfate deconjugation (ρ > 0.6, *p* < 0.05), including five individual arylsulfatases, two other sulfatases, and five sulfatase-maturation enzyme annotations. The discrepancy between individual and cumulative sulfatase gene associations may reflect the substantial functional diversity and substrate specificity of microbial sulfatases. Whereas individual sulfatases may contribute to TBBPA-sulfate hydrolysis, cumulative abundance measurements also include numerous sulfatases targeting unrelated substrates, potentially diluting associations with TBBPA deconjugation activity. Notably, several positively associated sulfatase and maturation-enzyme annotations were assigned to *Eubacterium rectale* and *Ruminococcus torques*, suggesting that specific microbial taxa may contribute to interindividual differences in TBBPA reactivation.

In contrast to the heterogeneous substrate specificities of sulfatases, anSMEs participate in the post-translational activation of multiple sulfatase substrates, potentially explaining why their collective abundance shows a stronger association with TBBPA-sulfate deconjugation. Many bacterial sulfatases require conversion of a conserved cysteine or serine residue into catalytically essential Cα-formylglycine (FGly), a modification catalyzed by sulfatase-maturation enzymes. Under anaerobic conditions, anSMEs utilize radical S-adenosylmethionine chemistry to generate FGly and activate their sulfatase substrates ^23–25, 31^. Consequently, the abundance of sulfatase-encoding genes alone may not accurately reflect the pool of catalytically active enzymes, particularly when maturation capacity varies among microbial communities. Moreover, bacterial sulfatases exhibit substantial substrate specificity, and many annotated sulfatases preferentially hydrolyze sulfated carbohydrates or other endogenous substrates rather than TBBPA-sulfate ^27, 32^. Thus, total sulfatase gene abundance may overestimate the community’s capacity to deconjugate TBBPA-sulfate. In contrast, anSME gene abundance may better reflect the potential capacity of the microbial community to generate catalytically competent sulfatases, providing a plausible explanation for its stronger association with measured TBBPA regeneration. These findings suggest that sulfatase maturation may represent an important functional constraint on microbial TBBPA-sulfate deconjugation, although direct enzyme activity measurements and targeted perturbation studies are needed to establish causality.

Taken together, the present Pilot-10 Human Donor Study provided initial evidence that human fecal microbiota can deconjugate TBBPA-monosulfate, regenerating unconjugated TBBPA with pronounced interindividual variability. The identification of putative fast, intermediate, and slow metabolizer phenotypes highlights the potential importance of individual gut microbial metabolic capacity in determining xenobiotic transformation. The absence of appreciable Tri-BBPA formation further suggests that microbial desulfation, but not downstream debromination, was the predominant transformation observed under our experimental conditions. Together with our previous findings on PCB-sulfate deconjugation, these results support microbial desulfation as a potentially broader and underrecognized mechanism of environmental xenobiotic reactivation.

Future studies will prioritize identifying the necessity of specific microbial taxa, sulfatases, and associated metabolic pathways responsible for TBBPA-sulfate deconjugation through integrated metagenomic, metatranscriptomic, and functional analyses. Mechanistic validation using targeted genetic disruption, antisense oligonucleotide-mediated gene suppression, and selective enzymatic inhibition will be essential to establish the necessity and substrate specificity of candidate microbial sulfatases. Expanding the investigation to larger, demographically diverse human cohorts will help validate the observed metabolizer phenotypes and determine whether microbial functional signatures can predict individual deconjugation capacity. Additionally, evaluating other TBBPA conjugates, including glucuronides, disulfates, and mixed conjugates, together with intestinal absorption models and in vivo toxicokinetic studies, will be important for establishing the broader consequences of microbial deconjugation for systemic exposure and toxicity.

## Acknowledgements

The authors would like to thank members of the Cui Laboratory for proof-reading the manuscript.

## Footnote

This work was supported by the National Institute of Environmental Health Sciences (NIEHS), National Institutes of Health (NIH), through grants R01ES031098, R01ES030197, and P30ES030287, Environmental Health and Microbiome Research Center (EHMBRACE) at University of Washington, as well as startup funding from the Department of Environmental and Molecular Toxicology, College of Agricultural Sciences, Oregon State University.

